# Gastric cancer derived exosomes induce peritoneal mesothelial cell EMT through TGF-β1/Smads pathway to promote peritoneal metastasis

**DOI:** 10.1101/2021.07.23.453534

**Authors:** Jungang Dong, Zhongbo Zhu, Guoning Cui, Zhixuan Zhang, Juan Yue, Yinghong Zhang, Xuehan Yao, Minfeng Huo, Jingjing Wei, Qingmiao Wang, Lirong Dai, Peiqing Li, Xi-Ping Liu

## Abstract

Epithelial-mesenchymal transition (EMT) plays an important role in peritoneal metastasis of Gastric cancer (GC). Tumor exosomes can mediate tumor directed metastasis, and TGF-β1 is an important factor in inducing tumor Epithelial mesenchymal transition. However, it is not clear whether GC derived exosomes can induce peritoneal mesothelial cells through the TGF-β1/ Smads pathway and the effect of injured peritoneal mesothelial cells on the biological characteristics of GC cells. In this study, we demonstrated that GC-derived exosomes can activate the TGF-β1/Smads pathway in peritoneal mesothelial cells and induce the corresponding EMT process, and that the injured peritoneal mesothelial cells can improve the migration and adhesion of GC cells. Taken together, these data further support the critical role of exosomes in the remodeling of the pre-metastatic microenvironment.

## Introduction

Gastric cancer (GC) is the second most common disease with cancer-related mortality in the world^[1,2]^, and this high mortality rate is mainly related to peritoneal metastasis caused by delayed diagnosis and treatment^[3,4]^. Peritoneal metastasis is common in advanced GC and usually leads to poor prognosis^[5]^. To date, little is known about the mechanism of peritoneal metastasis, so there is no effective treatment.

Peritoneal membrane is composed of monolayer mesothelial cells and subcutaneous connective tissue. Apoptosis and fibrosis of peritoneal mesothelial cells are more conducive to invasion of GC cells^[6]^, and intact peritoneal mesothelial cells can effectively resist such invasion^[7,8]^. It is well known that peritoneal metastasis is a complex process. According to the classical “seed soil” theory^[9]^, tumor cells form seeds out of the in-situ tumor, and the communication between the “seed” and the “soil” peritoneal membrane leads to the apoptosis of peritoneal mesothelial cells or the occurrence of EMT. The formation of a microenvironment conducive to the metastasis of cancer cells before settlement. It is inferred that the communication between the seed and the soil is a key link in the metastasis of cancer cells. As a bridge of communication between cells, exosomes may play an important role^[10]^.

It is reported that peritoneal mesothelial cells undergoing EMT is a marker of peritoneal metastasis^[11]^. Recent studies have found that exosomes derived from GC cells promote peritoneal metastasis by inducing EMT in peritoneal mesothelial cells through the MAPK/ERK pathway^[12]^. In addition, miR-21-5p and miR-106A in exosomes induce peritoneal metastasis of EMT in peritoneal mesothelial cells by targeting Smad7^[13,14]^. It is suggested that exosomes play an important role in the microenvironment before remodeling and metastasis. TGF-β1 is one of the key molecules in inducing epithelial-mesenchymal transformation of tumor cells^[15]^. However, it is not clear whether gastric exosomes can activate TGF-β1 / Smads pathway and induce EMT, in peritoneal mesothelial cells.

Based on the above findings, we hypothesized that before GC metastasis, the exosomes secreted by GC was absorbed by peritoneal mesothelial cells, and EMT, was induced by TGF-β1 / Smads pathway to promote peritoneal metastasis.

In this study, we isolated and purified GC derived exosomes, observed the uptake of GC derived exosomes by peritoneal mesothelial cells, detected the key molecules of TGF-β1/ Smads pathway and EMT-related markers, and examined whether peritoneal mesothelial cells affected the migration and adhesion of GC cells after EMT to verify our hypothesis.

### Materials and methods Cell lines and cultured

Human peritoneal mesothelial cells(HMrSV5,), MKN-45,BGC-823 and NCI-N87 were purchased from the cell bank of Shanghai Chinese Academy of Sciences. HMrSV5 were cultured in DMEM medium containing 10% high quality fetal bovine serum and 1% penicillin-streptomycin double antibody, and others cultured were cultured RPMI-1640 medium containing 10% high quality fetal bovine serum and 1% penicillin-streptomycin double antibody. They are all cultivated in 37 °C, 5% CO_2_ and saturated humidity incubator, the liquid was changed once every 3 days, and the cells were passaged at 80% fusion degree at 1:3.

### Reagents and antibodies

The specific TGF-β1/Smads inhibitor LY2109761 (HY-12075) was purchased from MCE, USA, Dil (41085-99-8) was purchased from Sigma, and DAPI (C1005) was purchased from Biyuntian, China for western blotting. The other antibodies were purchased from Abcam, UK.

### Purification and labeling of exosomes

MKN-45, BGC-823 and NCI-N87 cells are cultured under normal conditions. When the fusion degree reaches 80%-90%, the old culture medium is first absorbed from the culture bottle, washed with PBS for 2 times, added to the exosomes system to prepare the culture medium, and continued to be cultured for 72 h. 10ml of the cell culture supernatant was collected. The supernatant was collected by centrifugation 5min at 1500 × g at 4°C and centrifuged at 5000 × g at 4°C for 20min. The cell debris and other impurities were removed. Finally, the larger vesicles were removed by 0.22um filter membrane. The supernatant of the pretreated exosomes was centrifuged at 100, 000 ×g and centrifuged at 4 °C for 90min, and the supernatant was removed and PBS was added to re-suspension the exosomes to obtain the exosomes suspension. Exosome concentration was determined by BCA method and exosome marker proteins CD9 and CD63 were detected by Western Blot^[16]^. The obtained exosomes and Dil were mixed at a volume ratio of 1000:1 and placed in the dark for 30 minutes. Centrifuge for 90 min at 100,000xg, 4°C, resuspend in PBS and ultracentrifuge again to collect exosomes. It is GC derived exosomes labeled with DiI.

### Establishment of co-culture model

HMrSV5 cells from logarithmic growth phase were inoculated into cell culture plate at the density of 2 × 10^5^/ml. According to the experiment, the cells were divided into three groups: negative control group: HMrSV5 was cultured alone; model group: HMrSV5 was added with exosome labeled by 100ug/mL DiI; experimental group: culture medium with LY2109761 was added, and exosome labeled by 100ug/mL DiI was added at the same time; the culture was terminated after 72 h of culture.

### Western Blot

The cell samples were dissolved in the lytic buffer and phenylmethylsulfonyl fluoride(PMSF) on the ice. The supernatant was centrifuged at 12000 × g 4 °C for 15min, and the supernatant was added with 1 / 4 of 4 × sample buffer to shake evenly. The protein concentration was determined by boiling at 95 °C for 15 min. Then the lysate was decomposed by SDS-polypropylamide gel electrophoresis (SDS-PAGF). Transfer the gel to the Polyvinylidene fluoride (PVDF) membrane. Sealed with 5% skim milk powder, incubated overnight with primary antibody at 4 °C, and reacted with secondary antibody at room temperature for 1.5 h. Finally, the ECL exposure solution was mixed at 1:1 according to A: B solution, and then evenly covered on the whole film, the reaction 2mins was put into the exposure instrument to detect. Image J 8.0 software was used to measure the gray value and calculate the relative expression level of protein.

### Real-time Quantitative PCR

Total RNA was extracted from cells using Trizol reagent (Takara, Japan) according to the manufacturer’s instructions. RNA (1 µg) was reverse transcribed into cDNA by Reverse Transcription System (Vazyme, Nanjing, China). To quantify the mRNA levels of TGF-β1/Smads signaling pathway, real-time PCR was performed using SYBR Green Kit (Cwbio, Beijing, China) on a Bio-Rad CFX96 Detection System. The relative gene expression was normalized to GAPDH. Data were analyzed by using the comparative Ct method. The primers of target genes used in this study were listed in Supplementary **Table 1**.

Glyceraldehyde-3-phosphate dehydrogenase (GAPDH) was used to normalize RNA inputs and as endogenous normalization control. All assays were calculated on the basis of 2^-△△Ct.^

### Fluorescent labeling

The cells of each group were washed with 1 × PBS for 3 times, each time for 3 minutes, and 15min was fixed with 4% paraformaldehyde (prepared by PBS) at room temperature. After washing again, DAPI was added and incubated in the dark for 15 mins. The specimens were stained with nuclei, and the excess DAPI was washed by PBS. And use absorbent paper to dry the residual PBS on the climbing piece. The red fluorescence expression of HMrSV5 in each group was observed under laser confocal microscope after the tablets were sealed with anti-fluorescence quenching agent, and the MOD values of red fluorescence labeling of HMrSV5 in each group were analyzed with Image J 8.0 software.

### Cell invasion assays

GC cells were inoculated with 5 × 104/ml in the plate of transwell superior chamber, such as 50ul diluted Matrigel glue. According to the purpose of the experiment, HMrSV5 and HMrSV5+DiI-GC exudates were added into the lower chamber at a cell density of 2 × 10 ^5^ / ml. Incubated with 5%CO_2_ at 37 °C for 24 h. The cells left at the top of the membrane are wiped off with a cotton swab. The invaded cells were fixed in 4% paraformaldehyde and stained with crystal violet. The number of invaded cells was counted under microscope. There are at least three compound holes in each group.

### Cell adhesion assays

After trypsin digestion of HMrSV5 cells growing well in logarithmic phase, the cells were resuscitated with complete culture medium and the cell concentration was adjusted to 2 × 10^5^/mL cell suspension. The 96-well plate of 100 μL HMRSV5 cell suspension was absorbed, and there were 3 multiple holes in each group. The cells were cultured at 37 °C in 5%CO_2_ incubator. After adhering to the wall, the culture medium of each group was changed according to each experimental group and co-cultured for 72 hours. The MKN-45 cells which grew well in logarithmic phase were digested with trypsin and the cell precipitates were collected. After washing with PBS twice, add CFSE working solution to avoid light and incubate for 20 mins. After incubation, CFSE-labeled GC cell precipitates were collected by centrifugation and resuscitated with complete culture medium. The concentration of CFSE-GC cells was adjusted to 5 × 104 ml to avoid light. The main results were as follows: (1) 100ul CFSE-GC cell suspension was added to the peritoneal mesothelial cells co-cultured with exosomes for 72 hours and incubated at 37 °C in 5% CO_2_ incubator for 1 h. (2) after the end of culture, the culture medium was discarded and washed gently with PBS for 3 times to elute the unbound GC cells. (3) the cell adhesion was observed under the phase contrast microscope, and the fluorescence value (excitation wavelength 485 nm, emission wavelength 530nm) was read in the fluorescence enzyme meter.

### Statistical Analysis

Statistical analyses were carried out by GraphPad Prism Software (version 8). All values were presented as mean values ± SD. Two-way ANOVA for multiple groups and unpaired Student’s *t* test for two groups were applied for statistical analysis. P < 0.05 was considered statistically significant.

## Results

### HMrSV5 Uptake MKN45, BGC-823 and NCI-N87 cells exosomes

The exosomes were isolated and purified from the supernatant of GC cells. Under transmission electron microscopy, it was observed that exosomes derived from GC cells presented oval or round shape, with a complete double-layer envelopment structure, containing low-density substances, with particle sizes ranging from 40 to 60 nm, forming a typical “cup mouth” shape, as shown in the yellow arrow in the figure (Figure 1A). CD9 and CD63 markers of exosomes were significantly higher than those of GC cells by Western blot (P<0.05) (Figure 1B, C). In order to verify the uptake of GC derivd exosomes by HMrSV5, the exosomes were labeled with Dil dye and the red fluorescence expression was determined (Figure 2). Compared with the control group, it was found that the exosomes marked with red dye in the model group were absorbed by HMrSV5 and dispersed in the cytoplasm, and the MOD value was significantly increased (Table 2) (P<0.05). Compared with the model group of MKN45, the value of MOD in BGC-823, NCI-N87 group was significantly lower. Compared with the LY2109761 Group of MKN45, the value of MOD in BGC-823, NCI-N87 group was significantly higher. Based on the above results, we found that MKN45 exosomes are sensitive to peritoneal mesothelial cells and LY2109761 inhibitors. We confirmed that HMrSV5 uptake exosomes, and LY2109761 can block this process. Therefore, we choose MKN45 cells for follow-up experiments.

**TABLE 1.**
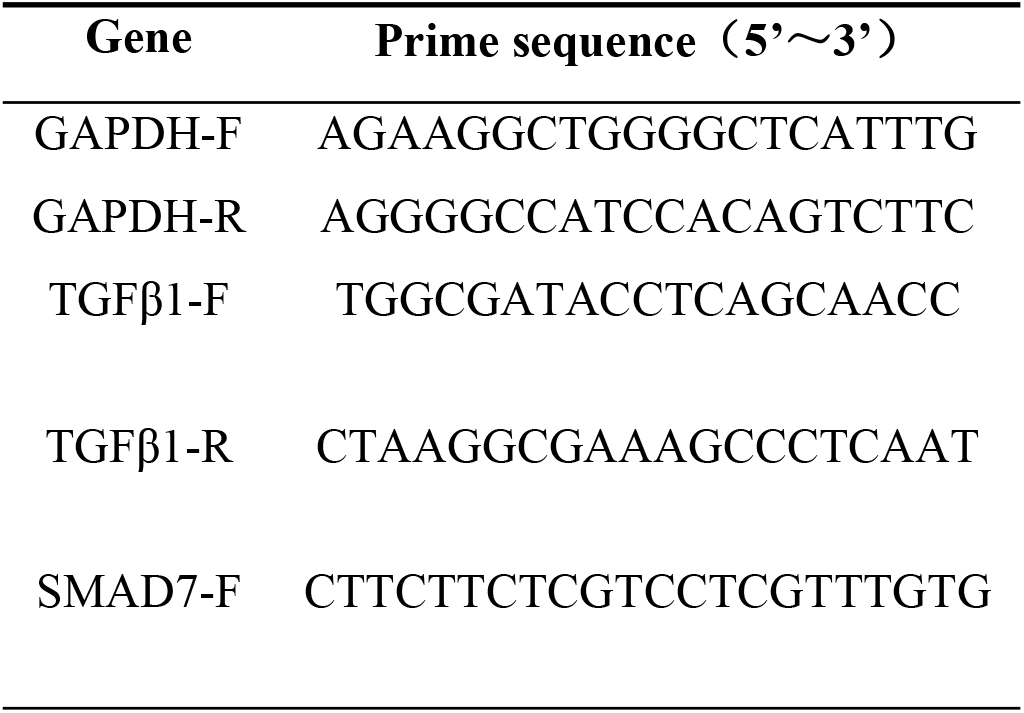
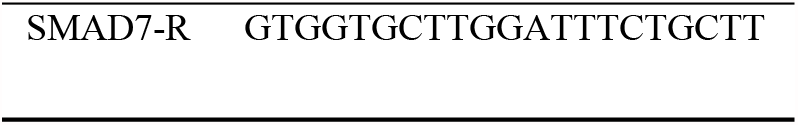
The primer sequence of a gene of GAPDH, TGF-β1 and Smad7.

**TABLE 2.** MOD values of gastric cancer exosomes were taken by HMrSV5 in each group (*n*=9). 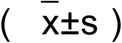

| Group | *MKN45* | **BGC-823** | **NCL-N87** |
| --- | --- | --- | --- |
| Blank | 0.00±0.00 | 0.00±0.00 | 0.00±0.00 |
| Model | 8492.95±99.38 | 8114.40±230.63* | 7737.96±196.31* |
| LY2109761 | 6835.97±362.13 | 7798.53±750.62<sup>#</sup> | 7639.63±469.34<sup>#</sup> |

Model Group :Compared with *MKN45* Group \*P < 0.05; LY2109761 Group :comparison with *MKN45* Group <sup>#</sup>P < 0.05

**FIGURE 1.**
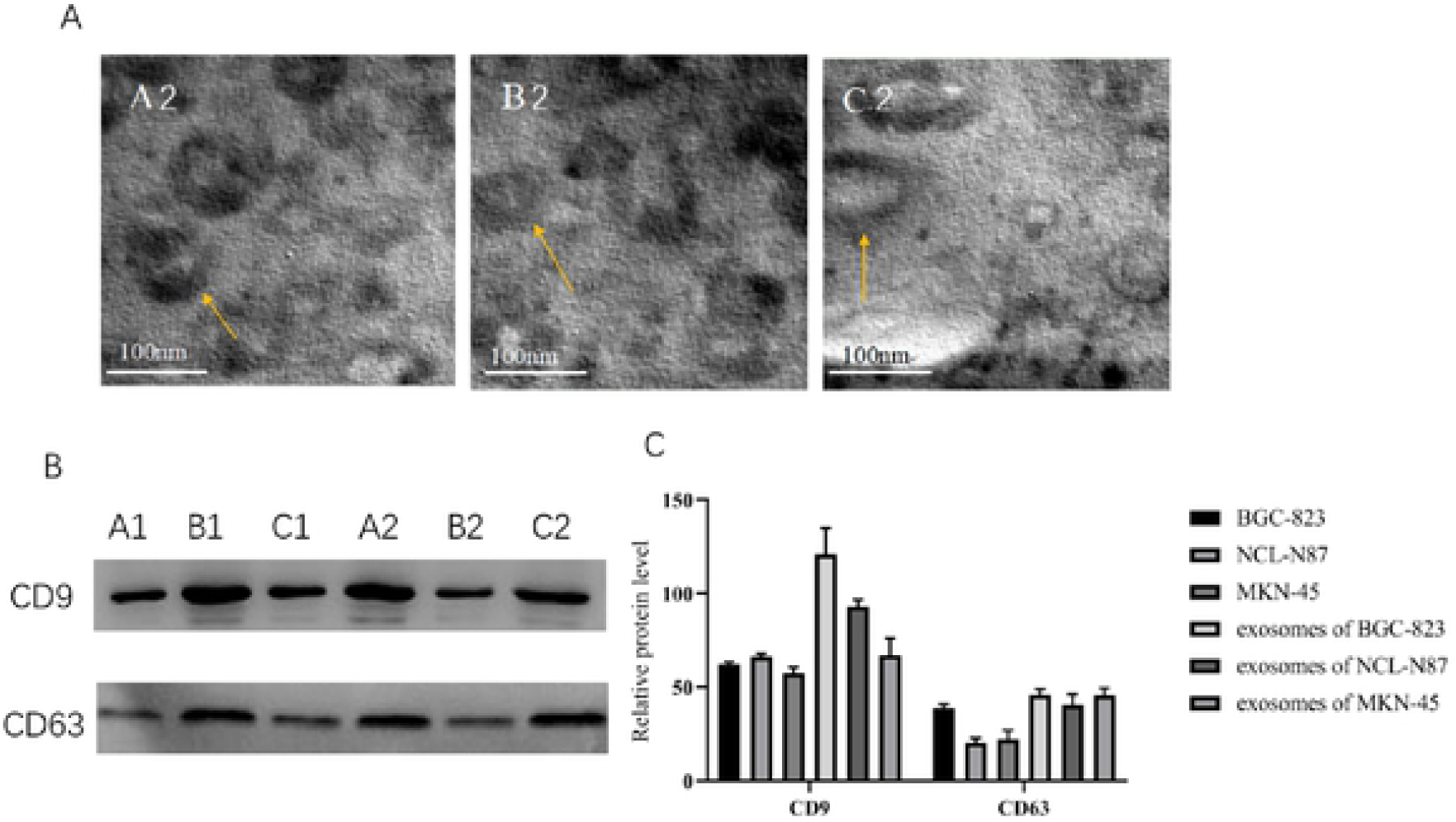
Identification and characterization of exosomes secreted by BGC-823, NCL-N87 and MKN-45cells A1, B1, C1 represent BGC-823, NCL-N87 and MKN-45 cells ; A2, B2, C2 represent the exosomes of BGC-823, NCL-N87 and MKN-45 cells (A) BGC-823, NCL-N87 and MKN-45cells exosomes were visualized by electron microscopy (×30,000).Scale bar = 100 nm. (B) Western blotting shows that BGC-823, NCL-N87 and MKN-45cells exosomes expressed exosome-associated protein, including CD9 and CD63. (C) CD9 and CD63 proteins content in BGC-823, NCL-N87 and MKN-45cells exosomes.

**FIGURE 2.**
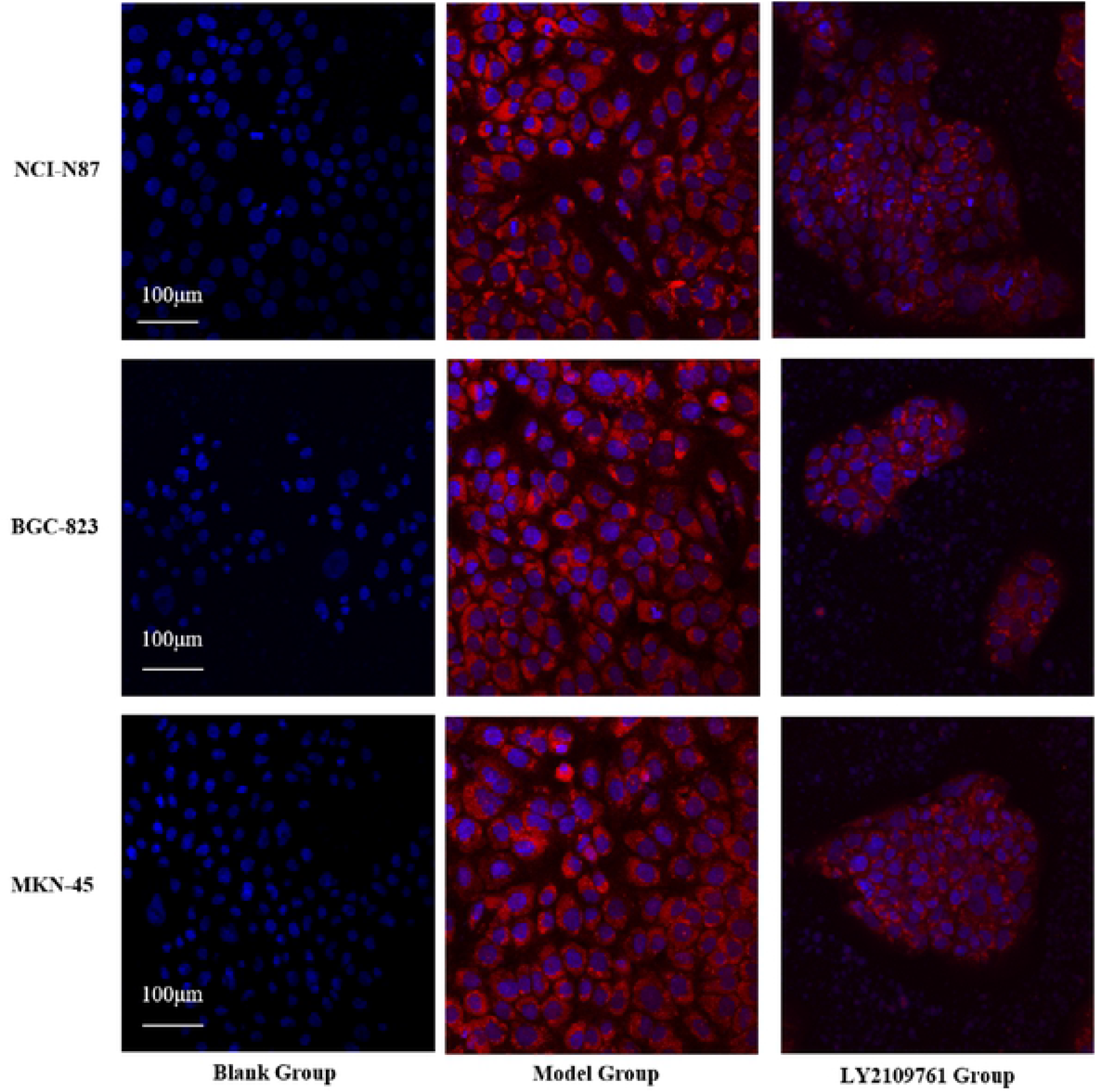
Uptake of BGC-823, NCL-N87 and MKN-45 cells exosomes by HMrSV5 after 72 hours of co-culture.

### Exosomes derived from GC induce EMT in HMrSV5 cells

In order to verify the mechanism that GC derived exosomes can induce EMT and EMT in HMrSV5 cells, GC derived exosomes were co-cultured with HMrSV5 and LY2109761 inhibitors were added at the same time. The effect of GC exosome on HMrSV5 cells was observed by inverted microscope **(Figure 3A)**. Compared with the control group, it was found that the morphology and intercellular junction of MKN-45 cells changed significantly after incubation with exosomes of HMrSV5 cells for 72 h. The HMrSV5 cells in the control group showed typical polygonal and pebble-like appearance and the cells were closely arranged. In the model group, HMrSV5 cells showed spindle-like appearance and loose arrangement, and the density increased. After the addition of LY2109761 inhibitors, the density decreased and the arrangement was more compact. In order to better observe the expression of EMT-related markers in HMrSV5 cells induced by gastric exosomes **(Figure 3B)**. Compared with the normal group, the expression of E-cadherin and CK19 protein decreased and the expression of α-SMA and Elastin protein increased in the model group (P<0.05), while the expression of E-cadherin and CK19 protein increased and the expression of α-SMA and Elastin protein decreased in the inhibitor group compared with the model group (P<0.05).

**FIGURE 3.**
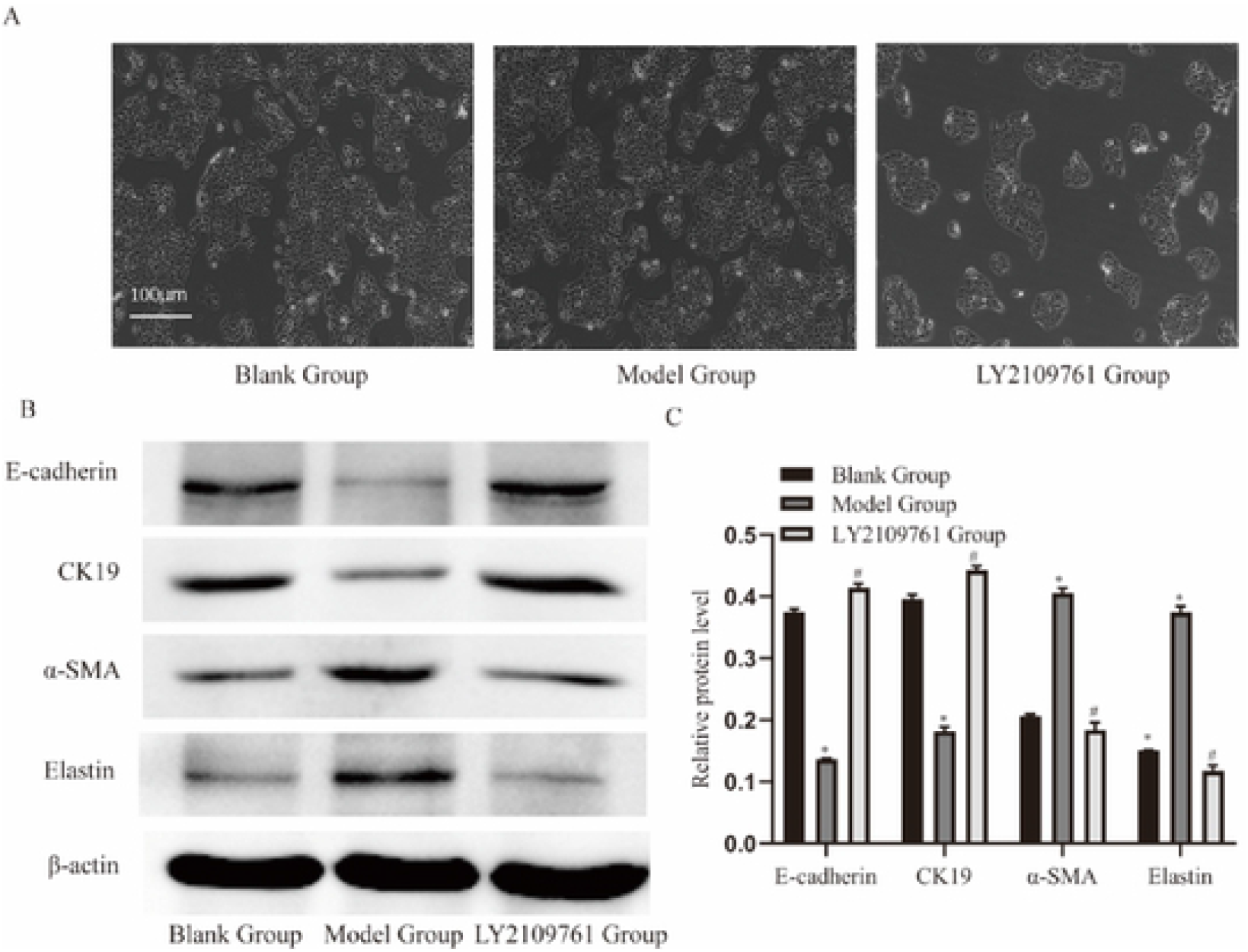
MKN-45cells exosomes foster the EMT of HMrSV5. **(A)** The effects of MKN-45 cells exosomes on HMrSV5 were assessed by Inverted microscope. **(B)** The expression of EMT related markers in HMrSV5 treated with MKN-45cells exosomes was determined by western blot **(C)** The quantification of protein expression level. Comparison with blank group. ^**\***^ P < 0.05, comparison with model group ^#^P <0.05.

### MKN-45 cell exosomes activate HMrSV5 through the TGF-β1/smads signalling pathway

In order to further study the mechanism of MKN-45 exosome-induced HMrSV5 secretion EMT, we detected the activation of TGF-β1 / Smads pathway (figure 3). It was observed that the levels of TGF-β1, p-smad2 and p-smad3 were significantly up-regulated after co-culture with MKN-45 exosomes, while the total levels of smad2 and smad3 remained similar **(Figure 4B)**. These results clearly showed that the TGF-β1 / Smads pathway was highly activated when HMrSV5 was co-cultured with MKN-45 exosomes. After LY2109761 treatment, TGF-β1 and P-Smad2/3 decreased significantly, while Smad7 increased significantly **(Figure 4C)**. The expression of TGF-β1 and Smad7 mRNA was detected by QPCR. The results showed that LY2109761 almost completely blocked the expression of TGF-β1 induced by exosome in MKN-45 cells and promoted the expression of Smad7 gene **(Figure 4A)**. This may contribute to the damage to the function and integrity of mesothelial cells in the prevention of gastric tumor invasion / metastasis. In addition, LY2109761 treatment inhibited the epithelial-mesenchymal transformation of HMrSV5 cells **(Figure 4C)**. Taken together, these results suggest that the exosomes of MKN-45 cells induce EMT in HMrSV5 cells by activating the TGF-β1 / Smads pathway. LY2109761, an inhibitor of TGF-β1 / Smads pathway, blocks this process.

**FIGURE 4.**
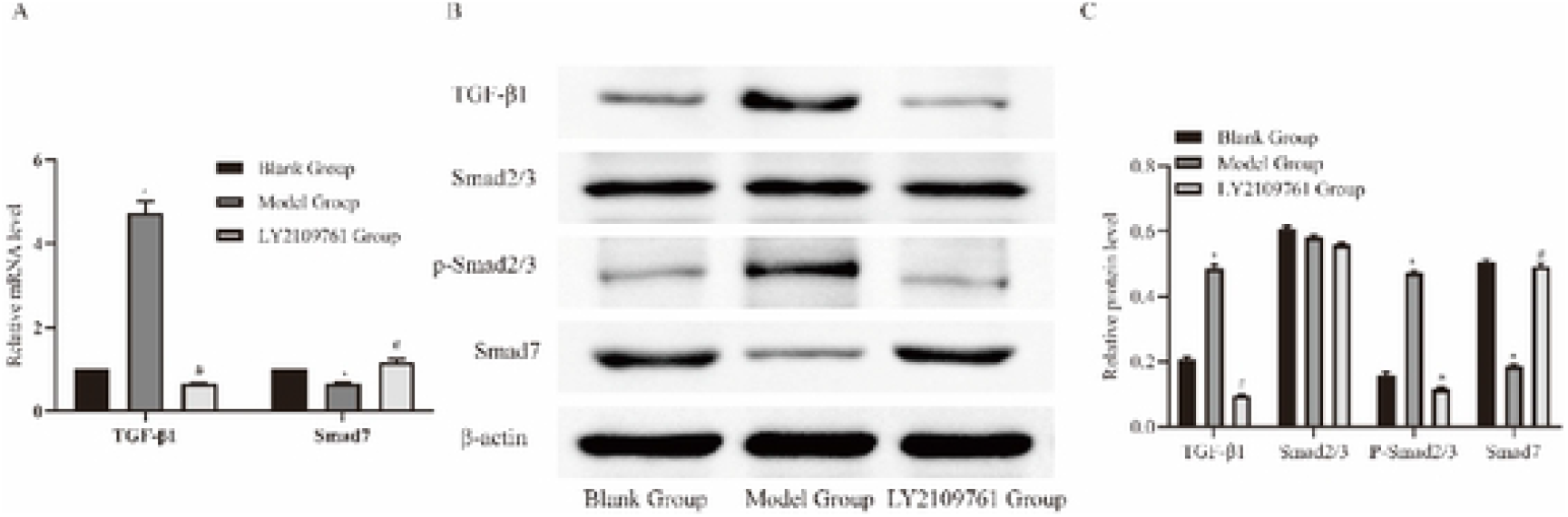
MKN-45 cells Exosomes activate TGF-β1/Smads signaling pathways in HMrSV5. **(A)** RT -PCR analyses of the expression of the cell signaling pathway in HMrSV5 treated with MKN-45 cells exosomes. Data shown above were the means ± SD of three independent experiments. **(B)** The expression of cell signaling pathway in HMrSV5 treated with MKN-45cells exosomes was determined by western blot **(C)** The quantification of protein expression level. Comparison with blank group. ^**\***^P <0.05, comparison with model group ^#^P <0.05.

### HMrSV5 cells after EMT promote MKN-45 cell invasion and adhesion

In order to observe the effect of injured peritoneal mesothelial cells on the biological characteristics of GC cells, the invasion and adhesion of GC cells were detected **(Figure 5)**. We found that compared with normal HMrSV5 cells and MKN-45 cells cultured alone, HMrSV5 cells cultured in gastric exosomes could significantly improve the invasion and adhesion ability of MKN-45 cells. After the addition of inhibitor LY2109761, the invasion and adhesion ability of MKN-45 cells decreased.

**FIGURE 5.**
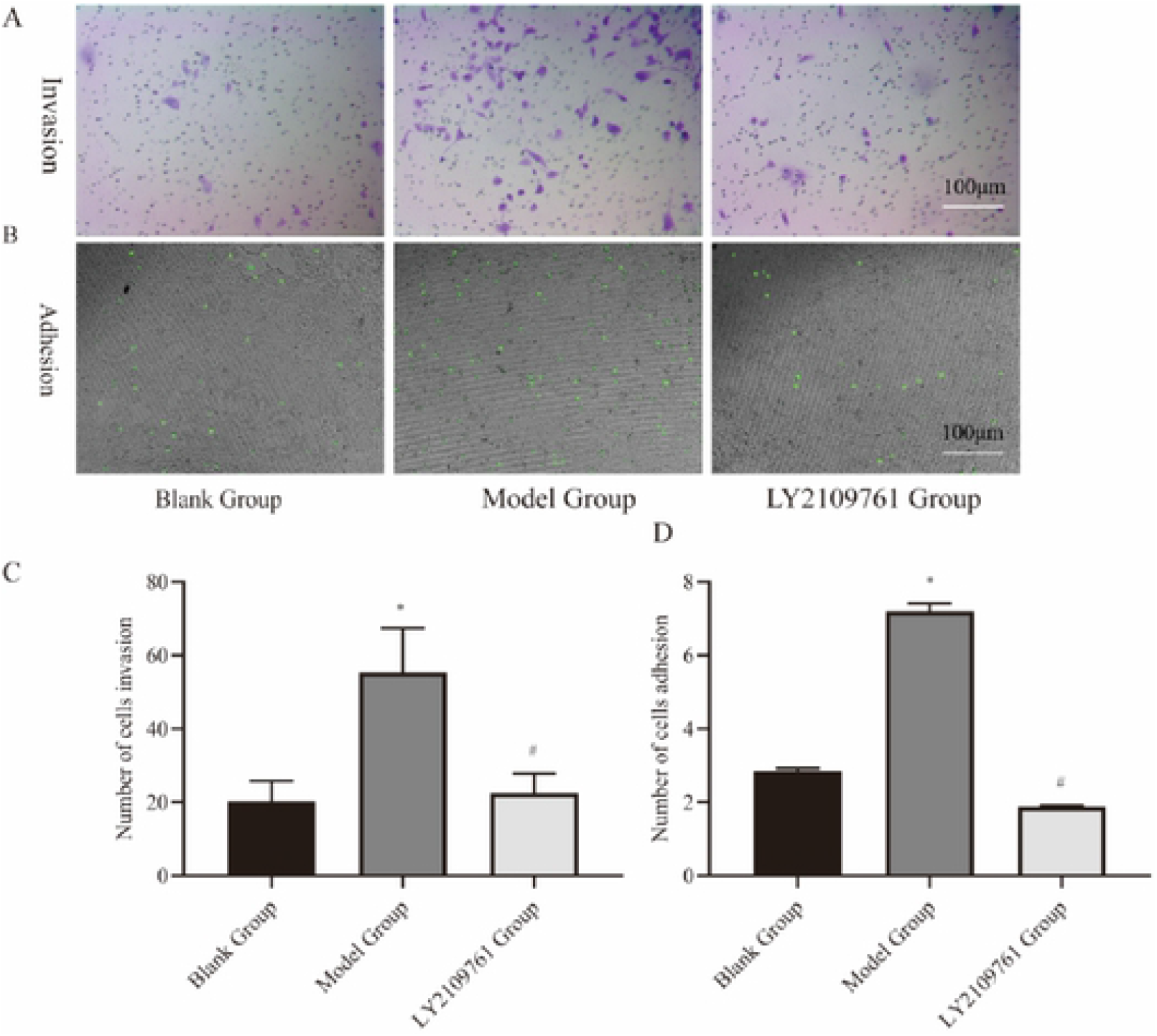
HMrSV5 cells after injury foster the invasion and adhesion of MKN-45 cells **(A)** Compare the invasion of MKN-45 cells by HMrSV5 cells after treatment with MKN-45 exosomes. **(B)** Compare the adhesion of MKN-45 cells by HMrSV5 cells after treatment with MKN-45 exosomes **(C)** The statistical analysis of the number of penetrated cells is shown in here. Scale bar: 100 µm. Data shown above were the means ± SD of three independent experiments. Compared with blank group ^**\***^P <0.05, comparison with model group #P < 0.05. **(D)** The statistical analysis of the fluorescence of adherent cells is shown in here. Data shown above were the means ± SD of three independent experiments. Compared with blank group ^**\***^ P < 0.05, comparison with model group #P< 0.05.

## Discussion

In this study, we observed that GC derived exosomes can induce HMrSV5 tumor EMT, to promote peritoneal metastasis. In order to elucidate the mechanism of GC exosomes on HMrSV5 secretion EMT, we interfered with LY2109761, an inhibitor of TGF-β1. At the same time, we detected the expression of TGF-β1, Smad2/3, P-Smad2/3 and Smad7, the key molecules of TGF-β1 / Samds signaling pathway, and the expression of E-cadherin, CK19, α-SMA and Elastin proteins in HMrSV5 cells, and observed the biological effects of HMrSV5 EMT on GC cells. Therefore, we can verify our previous hypothesis that GC exosomes induce EMT in peritoneal mesothelial cells through TGF-β1 / Smads pathway to promote peritoneal metastasis. At present, the mainstream view about peritoneal metastasis is the theory of cancer cell spread^[9,17]^, but some scholars believe that hematogenous metastasis is the main way of peritoneal metastasis^[18]^. Therefore, the mechanism of peritoneal metastasis remains to be fully elucidated, and this experiment is designed based on the theory of dissemination. In the past few years, the formation of metastatic microenvironment by exosomes before tumor metastasis has attracted a lot of attention, and it is considered to be the main messenger for long-distance transport of biological information in cancer cells. As previously reported, exosomes guide targeted metastasis of cancer cells through the microenvironment before reconstruction and metastasis of various cancers, such as renal cancer, GC, melanoma, pancreatic cancer, and breast cancer^[19-23]^. In this study, we observed that the exosomes of MKN-45 cells were absorbed by HMrSV5 cells, resulting in down-regulation of E-cadherin and CK19 proteins and up-regulation of α-SMA and Elastin proteins in HMrSV5 cells. Combined with the study of Deng, down-regulation of Zo-1 and up-regulation of vimentin^[12]^ fully proved that the occurrence of EMT, in HMrSV5 cells provided a favorable environment for GC cells to metastasize to the peritoneum. The above observations further indicate that exosomes play an important role in the construction of pre-metastatic microenvironment.

TGF-β1 has been reported to be an important factor causing EMT in peritoneal mesothelial cells^[15,24-26]^. In addition, EMT mediated by TGF-β1 may be important for the formation of pre-metastatic microenvironment in peritoneal metastasis^[27]^. It is worth noting that cancer exosomes bodies contain TGF-β1, which can play a biological function ^[28]^. Our results show that the presence of TGF-β1 in the exosomes of GC is the cause of peritoneal mesothelial cell injury.

Our significant finding is the activation of TGF-β1 / Smads pathway after peritoneal mesothelial cell injury induced by exosomes. TGF-β1 / Smads pathway is widely involved in the regulation of cell proliferation, differentiation, death and migration^[29]^. In addition, it has been previously reported that TGF-β1 can cause EMT in peritoneal mesothelial cells. In this study, different from the Deng study that GC exosome activated MAPK/ERK pathway to induce peritoneal mesothelial cell injury, we proved that GC exosome can also activate TGF-β1 / Smads pathway to induce peritoneal mesothelial cell injury. Administration of LY2109761, a specific TGF-β1 inhibitor, successfully reversed the EMT process of peritoneal mesothelial cells.

Our next significant finding is that EMT in peritoneal mesothelial cells improves the migration and adhesion ability of GC cells. Different from previous studies, we not only proved the injury of interperitoneal cells induced by GC exosomes, but also verified the effect of injured peritoneal mesothelial cells on the biological characteristics of GC cells. Interestingly, normal peritoneal mesothelial cells can also induce the migration and adhesion of GC cells, which seems to have been confirmed by OlgaKersy et al. OlgaKersy[30] study found that healthy peritoneal tissues of patients with GC without peritoneal spread can also promote the growth, migration and invasion of GC cells. It is further suggested that there may be two-way communication between GC cells and peritoneal mesothelial cells in the process of peritoneal metastasis of GC, which may enhance the metastasis of GC to the peritoneum, which needs to be further verified by experiments.

In conclusion, this study shows that GC exosomes can activate the TGF-β1 / Smads pathway of peritoneal mesothelial cells, and the peritoneal mesothelial cells after EMT, damage can further promote the migration and adhesion of GC, thus promoting the peritoneal metastasis of GC.

## Author contributions

Jungang Dong and Zhongbo Zhu wrote the manuscript and cultured the cells and tested them. Xiping Liu and Peiqing Li generated the conception and design of the study. Guoning Cui helped perform the experiments and collect data. Other authors helped perform the analysis with constructive discussions. All authors read and approved the final manuscript.

## Acknowledgments

This work was supported by the National Natural Science Foundation of China (81860813, 81860782) and Longyuan Youth Innovation and Entrepreneurship Talent Project (2020RCXM183).

---

**TABLE 1** The primer sequence of a gene of GAPDH, TGF-β1 and Smad7.

**TABLE 2** MOD values of gastric cancer exosomes were taken by HMrSV5 in each group (*n*=9).

